# The ZuCo Benchmark on Cross-Subject Reading Task Classification with EEG and Eye-Tracking Data

**DOI:** 10.1101/2022.03.08.483414

**Authors:** Nora Hollenstein, Marius Tröndle, Martyna Plomecka, Samuel Kiegeland, Yilmazcan Özyurt, Lena A. Jäger, Nicolas Langer

## Abstract

We present a new machine learning benchmark for reading task classification with the goal of advancing EEG and eye-tracking research at the intersection between computational language processing and cognitive neuroscience. The benchmark task consists of a cross-subject classification to distinguish between two reading paradigms: normal reading and task-specific reading. The data for the benchmark is based on the Zurich Cognitive Language Processing Corpus (ZuCo 2.0), which provides simultaneous eye-tracking and EEG signals from natural reading. The training dataset is publicly available, and we present a newly recorded hidden testset. We provide multiple solid baseline methods for this task and discuss future improvements. We release our code and provide an easy-to-use interface to evaluate new approaches with an accompanying public leaderboard: www.zuco-benchmark.com.

**Highlights:**

- We present a new machine learning benchmark for reading task classification with the goal of advancing EEG and eye-tracking research.
- We provide an interface to evaluate new approaches with an accompanying public leaderboard.
- The benchmark task consists of a cross-subject classification to distinguish between two reading paradigms: normal reading and task-specific reading.
- The data is based on the Zurich Cognitive Language Processing Corpus of simultaneous eye-tracking and EEG signals from natural reading.

## 1. Introduction

Reading plays a fundamental role in the acquisition of information (e.g., encyclopedias) and communication (e.g., emails). As we read, our eyes gaze through the written sentences in a sequence of fixations and high-velocity saccades to extract visual information which are forwarded to the brain to obtain meaning. Thus, assessing where a person looks during reading while recording brain activity non-invasively with electroencephalography (EEG) provides powerful behavioral and physiological measures for cognitive neuroscience to further the understanding of human language processing. Most previous experimental reading research has used hand-picked reading materials in highly controlled experimental settings (Brennan, 2016; Nastase et al., 2020). The neural correlates of reading have traditionally been studied with serial word-by-word presentation with a fixed presentation time, which eliminates important aspects of the natural reading process and precludes direct comparisons between neural activity and oculomotor behavior (Dimigen et al., 2011; Kliegl et al., 2012). The electrical neural correlates of normal reading of naturally occurring real-world sentences have been investigated less frequently due to a number of methodological challenges related to identifying the exact timing and type of visual stimuli presented during reading.

Because of recent methodological progress in stimulus presentation and data preprocessing (Dimigen et al., 2011; Ehinger and Dimigen, 2019), an excellent temporal resolution, and low costs, co-registered EEG and eye-tracking have become important tools for studying the temporal dynamics of naturalistic reading (Frey et al., 2018; Hollenstein et al., 2018). Fixation-related potentials (FRPs), the evoked electrical responses time-locked to the onset of fixations, have become important tools for researchers to study various topics including free-viewing visual perception (e.g., Rämä and Baccino, 2010), brain-computer interfaces (e.g., Finke et al., 2016), and natural reading (e.g., Degno et al., 2019). In naturalistic reading paradigms, FRPs allow the study of the neural dynamics of how new information from a currently fixated word affects the ongoing language comprehension process.

In this work, we leverage these novel methodological advances to offer a machine learning (ML) benchmark challenge, formulated as a cross-subject classification task, to identify two reading tasks as accurately as possible. Specifically, the challenge is to discriminate between normal reading (with the only task of reading comprehension) and task-specific reading (with the purpose of finding specific information in the text) from eye-tracking and EEG data. Decoding mental states and detecting specific cognitive processes occurring in the brain during different reading tasks (i.e., *reading task classification*) are important challenges in cognitive neuroscience as well as in natural language processing (NLP). Applications of reading task classification include measuring attention and engagement (Miller, 2015; Abdelrahman et al., 2019), detecting proper reading versus skimming (Biedert et al., 2012), as well as applications related to intent recognition within brain computer interfaces (Schalk et al., 2008). Other studies have demonstrated that recognizing reading patterns for estimating reading effort can improve the diagnosis of reading impairments such as dyslexia (Rello and Ballesteros, 2015; Raatikainen et al., 2021) and attention deficit disorder (Tor et al., 2021). Furthermore, it has been shown that using EEG and eye-tracking signals facilitates the prediction workload (Lobo et al., 2016) and investigation of language learning (Notaro and Diamond, 2018).

The accurate distinction of the cognitive processes occurring in different reading tasks is also important for ML and NLP. Identifying specific reading patterns can improve models of human reading and provide insights into human language understanding and how we perform linguistic tasks. This knowledge can then be applied to ML algorithms for NLP (e.g., information extraction applications). Computational models of language understanding can be adapted based on the insights from different reading and language processing tasks. Therefore, the identification of reading intents can be beneficial for computational methods of language understanding, but also for applications such as digital assistant tools, e.g., supporting translation processes, understanding how learners approach tasks in adaptive e-learning, and inferring document relevance.

A crucial potential of human physiological data in the context of NLP is that it can be leveraged to understand and to improve the manual labelling process required for generating training samples for supervised ML. For instance, Tokunaga et al. (2017) analyze eye-tracking data during the annotation of text to find effective gaze features for a specific NLP task and Tomanek et al. (2010) build cost models for active learning scenarios based on insights from eye-tracking data.

Reading task classification can help to improve the labelling processes by detecting tiredness from brain activity data and eye-tracking data, and subsequently to suggest breaks or task switching, or by using cognitive data directly to (pre-)annotate samples used for training ML models. If we can find and extract the relevant aspects of text understanding and annotation directly from the source, i.e., eye-tracking and brain activity signals during reading, we can potentially replace this expensive manual labelling work with ML models trained on physiological activity data recorded from humans while reading. Therefore, successful reading task classification could support the reduction of manual labor, improving label quality in ML systems as well as the job quality of annotators.

Essential for using neurophysiological signals to advance NLP is the availability of a large dataset providing concurrent measures of eye-tracking and EEG data, as well as ground truth labels for ML tasks. For the present benchmark, this is possible by leveraging a naturalistic dataset of reading English sentences, the Zurich Cognitive Language Processing Corpus (Hollenstein et al., 2018, 2020). The ZuCo dataset is publicly available and has recently been used in a variety of applications including leveraging EEG and eye-tracking data to improve NLP tasks (Barrett et al., 2018; Mathias et al., 2020; McGuire and Tomuro, 2021), evaluating the cognitive plausibility of computational language models (Hollenstein et al., 2019b; Hollenstein and Beinborn, 2021), investigating the neural dynamics of reading (Pfeiffer et al., 2020), developing models of human reading (Bautista and Naval, 2020; Bestgen, 2021). Recently, ZuCo has also been leveraged for an ML competition on eye-tracking prediction (Hollenstein et al., 2021a). This shows that the ZuCo dataset has been used successfully for a wide range of ML tasks.

To conclude, the contributions of our work can be summarized as follows: First, we formulate a benchmark task for applying ML techniques to an important problem in cognitive science, namely, the classification of cognitive tasks. Second, we provide the data^1^ and code^2^ to reproduce our experiments. We provide a public benchmark and leaderboard on a new held-out test data. All information can be found here: www.zuco-benchmark.com. Finally, we propose and discuss models using various feature sets as baseline models for this benchmark task. We present detailed analyses of the results for both eye-tracking and EEG features and discuss the model performances.

## 2. Methods

The basis for this ML benchmark task is the Zurich Cognitive Language Processing Corpus 2.0 (ZuCo 2.0). ZuCo 2.0 was originally published in Hollenstein et al. (2020). In short, this corpus contains gaze and brain activity data of 18 participants reading 739 English sentences, 349 in a normal reading paradigm, and 390 in a task-specific paradigm, in which the participants actively search for a semantic relation type in the given sentence as a linguistic annotation task. This new dataset provides experiments designed to analyze the differences in cognitive processing between normal reading and task-specific reading.

In previous work, we recorded a first dataset (i.e., ZuCo 1.0) of simultaneous eye-tracking and EEG during natural reading (Hollenstein et al., 2018). ZuCo 1.0^3^ consists of three reading tasks, two of which contain very similar reading material and experiments as presented in the current work. However, for ZuCo 1.0 the normal reading and task-specific reading paradigms were recorded in different sessions on different days. Therefore, the recorded data from ZuCo 1.0 is not appropriate as a means of comparison between normal reading and task-specific reading, since the differences in the brain activity data might result mostly from the different sessions due to the sensitivity of EEG. Therefore, while the data is available in the same format, it is not recommended to be used for this benchmark task. In the following section, we describe the compilation of the ZuCo 2.0 dataset.

### 2.1. Reading Materials

During the recording session, the participants read a total of 739 sentences that were selected from the Wikipedia corpus provided by (Culotta et al., 2006). This corpus was chosen because it provides annotations of semantic relations. Relation detection is a high-level semantic language understanding task requiring complex cognitive processing. ZuCo 2.0 includes seven of the originally defined relation types: *politicaL_affiliation, education, founder, wife/husband, job_title, nationality,* and *employer*. The sentences were chosen with similar sentence lengths and Flesch reading ease scores (Kincaid et al., 1975) between the two reading tasks. The dataset statistics are shown in Table 1.

**Table 1:** Descriptive statistics of reading materials (SD = standard deviation), including Flesch readibility scores.

|  | NR | TSR |
| --- | --- | --- |
| sentences | 349 | 390 |
| sent. length | mean (SD), range<br>19.6 (8.8), 5-53 | mean (SD), range<br>21.3 (9.5), 5-53 |
| total words | 6828 | 8310 |
| word types | 2412 | 2437 |
| word length | mean (SD), range<br>4.9 (2.7), 1-29 | mean (SD), range<br>4.9 (2.7), 1-21 |
| Flesch score | 55.38 | 50.76 |

Of the 739 sentences, the participants read 349 sentences in a normal reading paradigm and 390 sentences in a task-specific reading paradigm, in which they had to determine whether a certain relation type occurred in the sentence or not. Table 2 shows the distribution of the different relation types in the sentences of the task-specific annotation paradigm. Purposefully, there are 63 duplicates between the normal reading and the task-specific sentences (8% of all sentences). The intention of these duplicate sentences is to provide a set of sentences read twice by all participants with a different task in mind. Hence, this enables the comparison of eye-tracking and brain activity data when reading normally and when annotating specific relations. During both tasks, the participants were able to read in their own speed, using a control pad to move to the next sentence and to answer the control questions, which allowed for natural reading. Since all subject read at their own personal pace, the reading speed between varies between subjects. Figure 1 shows the average sentence length, reading speed, and omission rate for each task. The sentence length (i.e., the number of words per sentence) was controlled in the selection of reading materials, so that it would not differ significantly between the two tasks (NR mean 19.6, std 8.8; TSR mean 21.3, std 9.5).

**Figure 1:**
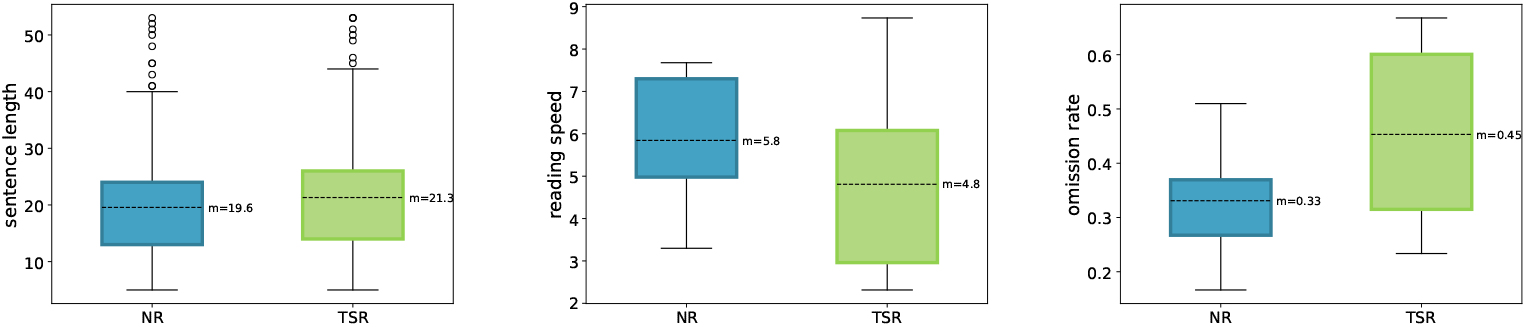
Sentence length (words per sentence), reading speed (seconds per sentence) and omission rate (percentage of words not fixated) comparison between normal reading (NR) and task-specific reading (TSR) of the sentence in ZuCo 2.0.

**Table 2:**
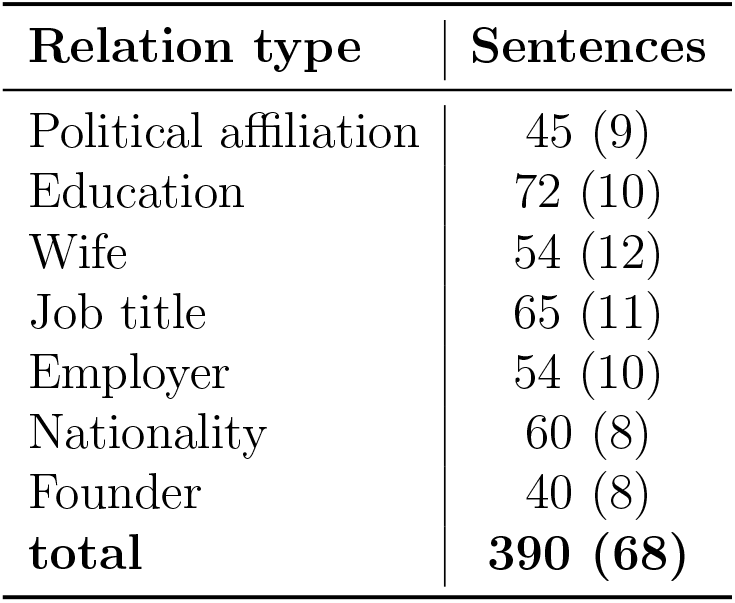
Distribution of relation types in the task-specific reading. The right column contains the number of sentences, and the number control sentences without a relation in brackets.

| Relation type | Sentences |
| --- | --- |
| Political affiliation | 45 (9) |
| Education | 72 (10) |
| Wife | 54 (12) |
| Job title | 65 (11) |
| Employer | 54 (10) |
| Nationality | 60 (8) |
| Founder | 40 (8) |
| **total** | **390 (68)** |

#### 2.1.1. Normal Reading (NR)

In the first task, participants were instructed to read the sentences naturally, without any specific task other than comprehension. An example sentence is “He served in the United Stated Army in World War II, then got a law degree from Tulane University”. The control condition for this task consisted of single-choice questions about the content of the previous sentence. 12% of randomly selected sentences were followed by a comprehension question with three answer options on a new screen, for example, “Which university did he get his degree from? (1) Austin University, (2) Tulane University, (3) Louisiana State University”.

#### 2.1.2. Task-specific Reading (TSR)

In the second task, the participants were instructed to search for a specific semantic relation in each sentence they read. Instead of comprehension questions, the participants had to decide for each sentence whether it contains the relation or not, i.e., they were actively annotating each sentence. An example sentence containing the relation *founder* is “After this initial success, Ford left Edison Illuminating and, with other investors, formed the Detroit Automobile Company”. 17% of the sentences did not include the particular relation type and were used as control conditions. All sentences within one recording block involved the same relation type. Each block was preceded by a short practice round, which described the relation type and was followed by three sample sentences, so that the participants would be familiar with the respective relation type.

### 2.2. Linguistic Assessment

As a linguistic assessment, the vocabulary and language proficiency of the participants was tested with the LexTALE test (Lexical Test for Advanced Learners of English, Lemhöfer and Broersma, 2012). This is an unspeeded lexical decision task designed for intermediate to highly proficient language users. The average LexTALE score over all participants was 88.54%. Moreover, we also report the scores the participants achieved with their answers to the reading comprehension control questions and their relation annotations. The detailed scores for all participants are also presented in Table 3.

**Table 3:**
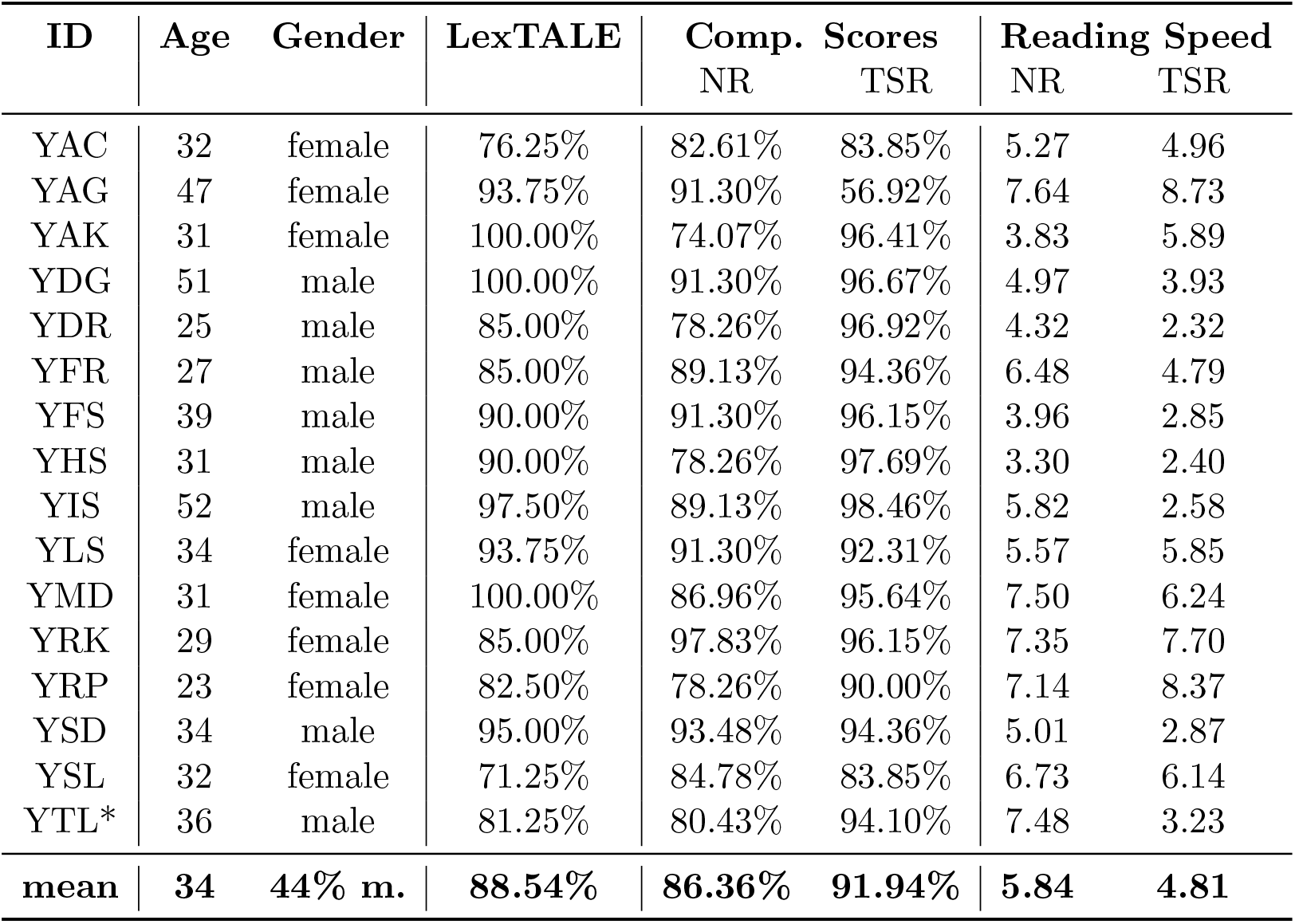
Subject demographics for ZuCo 2.0, LexTALE scores, scores of the comprehension questions, and individual reading speed (i.e., seconds per sentence) for each task. The * next to the subject ID marks a bilingual subject.

| ID | Age | Gender | LexTALE | Comp. Scores |  | Reading Speed |  |
| --- | --- | --- | --- | --- | --- | --- | --- |
|  |  |  |  | NR | TSR | NR | TSR |
| YAC | 32 | female | 76.25% | 82.61% | 83.85% | 5.27 | 4.96 |
| YAG | 47 | female | 93.75% | 91.30% | 56.92% | 7.64 | 8.73 |
| YAK | 31 | female | 100.00% | 74.07% | 96.41% | 3.83 | 5.89 |
| YDG | 51 | male | 100.00% | 91.30% | 96.67% | 4.97 | 3.93 |
| YDR | 25 | male | 85.00% | 78.26% | 96.92% | 4.32 | 2.32 |
| YFR | 27 | male | 85.00% | 89.13% | 94.36% | 6.48 | 4.79 |
| YFS | 39 | male | 90.00% | 91.30% | 96.15% | 3.96 | 2.85 |
| YHS | 31 | male | 90.00% | 78.26% | 97.69% | 3.30 | 2.40 |
| YIS | 52 | male | 97.50% | 89.13% | 98.46% | 5.82 | 2.58 |
| YLS | 34 | female | 93.75% | 91.30% | 92.31% | 5.57 | 5.85 |
| YMD | 31 | female | 100.00% | 86.96% | 95.64% | 7.50 | 6.24 |
| YRK | 29 | female | 85.00% | 97.83% | 96.15% | 7.35 | 7.70 |
| YRP | 23 | female | 82.50% | 78.26% | 90.00% | 7.14 | 8.37 |
| YSD | 34 | male | 95.00% | 93.48% | 94.36% | 5.01 | 2.87 |
| YSL | 32 | female | 71.25% | 84.78% | 83.85% | 6.73 | 6.14 |
| YTL* | 36 | male | 81.25% | 80.43% | 94.10% | 7.48 | 3.23 |
| **mean** | **34** | **44% m.** | **88.54%** | **86.36%** | **91.94%** | **5.84** | **4.81** |

### 2.3. Participants

The subjects from ZuCo 2.0 are provided as training data for the current benchmark. For the ZuCo 2.0, we recorded data from 19 participants and discarded the data of one of them due to technical difficulties with the eyetracking calibration. Another two subjects were discarded during data cleaning and preprocessing. Thus, we share the data of these 16 participants. All participants are healthy adults (between 23 and 52 years old; 10 females). Details on subject demographics can be found in Table 3. Their native language is English, originating from Australia, Canada, UK, USA or South Africa. Two participants are left-handed and three participants wear glasses for reading. All participants gave written consent for their participation and the re-use of the data prior to the start of the experiments. The study was conducted under approval by the Ethics Commission of the University of Zurich.

#### ZuCo 2.0 Held-out Testset

To provide a true hidden dataset for the current benchmark, we recorded data from 10 additional participants (i.e., a held-out testset). They underwent the identical procedure as in the ZuCo 2.0 dataset. All participants are healthy adults (mean age = 31.8 (SD=5.11), 4 females). All participants are right-handed. Their native language is English, originating from UK, Canada or USA. For an overview on subjects demographics, comprehension scores and reading speed please refer to Table 4. All participants gave written consent for their participation and the re-use of the data prior to the start of the experiments.

**Table 4:** Subject demographics for the new held-out test dataset, LexTALE scores, scores of the comprehension questions, and individual reading speed (i.e., seconds per sentence) for each task

| ID | Age | Gender | LexTALE | Comp. Scores |  | Reading Speed |  |
| --- | --- | --- | --- | --- | --- | --- | --- |
|  |  |  |  | NR | TSR | NR | TSR |
| XAH | 25 | female | 95.25% | 91.30% | 93.58% | 5.58 | 3.94 |
| XBB | 37 | male | 95.75% | 82.60% | 93.84% | 6.88 | 5.67 |
| XBD | 32 | male | 89.00% | 91.30% | 96.15% | 7.31 | 4.48 |
| XDT | 25 | male | 97.50% | 86.95% | 93.85% | 8.24 | 8.54 |
| XLS | 28 | male | 85.00% | 89.13% | 94.87% | 7.52 | 5.68 |
| XPB | 29 | male | 97.50% | 86.95% | 91.02% | 7.87 | 6.53 |
| XSE | 31 | female | 90.00% | 89.13% | 96.15% | 7.23 | 3.75 |
| XSS | 42 | female | 97.50% | 89.13% | 96.67% | 7.49 | 6.21 |
| XTR | 34 | female | 93.75% | 89.13% | 96.15% | 9.18 | 5.91 |
| XWS | 35 | male | 100.00% | 89.13% | 95.64% | 6.65 | 4.29 |
| **mean** | **31.8** | **60% m.** | **94.13%** | **88.48%** | **94.79%** | **7.40** | **5.50** |

### 2.4. Procedure

Data acquisition took place in a sound-attenuated and dark experiment room. Participants were seated at a distance of 68cm from a 24-inch monitor (ASUS ROG, Swift PG248Q, display dimensions 531×299 mm, resolution 800×600 pixels resulting in a display: 400×298.9 mm, a vertical refresh rate of 100 Hz). All sentences were presented at the same position on the screen and could span multiple lines. The sentences were presented in black on a light grey background with font size 20-point Arial, resulting in a letter height of 0.8 mm. The experiment was programmed in MATLAB 2016b (MATLAB, MathWorks, 2000), using PsychToolbox (Brainard, 1997). A stable head position was ensured via a chin rest. Participants were instructed to stay as still as possible during the recordings to avoid motor EEG artifacts. Participants completed the tasks sitting alone in the room, while two research assistants were monitoring their progress in the adjoining room. All recording scripts including detailed participant instructions are available alongside the data. During both tasks, the participants were able to read in their own speed, using a control pad to move to the next sentence and to answer the control questions, which allowed for natural reading. All 739 sentences were recorded in a single session for each participant. The duration of the recording sessions was between 100 and 180 minutes, depending on the time required to set up and calibrate the devices, and the personal reading speed of the participants. Participants were also offered snacks and water during the breaks and were encouraged to rest. We recorded 14 blocks of approx. 50 sentences, alternating between tasks: 50 sentences of normal reading, followed by 50 sentences of task-specific reading. The order of blocks and sentences within blocks was identical for all subjects. Each sentence block was preceded by a practice round of three sentences and followed by a short break to ensure a clear separation between the reading tasks. For the held-out test dataset, all blocks were merged and the order of the sentences was shuffled, before sharing the data on OSF, to prohibit the possibility that challenge participants would simply train a model to identify a block than the type of reading for each sentence.

### 2.5. Data Acquisition

#### 2.5.1. Eye-Tracking Acquisition

Eye movements and pupil size were recorded with an infrared video-based eye tracker (EyeLink 1000 Plus, SR Research) at a sampling rate of 500 Hz and an instrumental spatial resolution of 0.01°. The eye tracker was calibrated with a 9-point grid at the beginning of the session and re-validated before each block of sentences. Participants were instructed to keep their gaze on a given point until it disappeared. If the average error of all points (calibration vs. validation) was below 1° of visual angle, the positions were accepted. Otherwise, the calibration was redone until this criterion was reached.

#### 2.5.2. EEG Acquisition

We recorded the high-density EEG data at a sampling rate of 500 Hz with a bandpass of 0.1 to 100 Hz, using a 128-channel EEG Geodesic Hydrocel system (Electrical Geodesics). The *Cz* electrode served as a recording reference. The impedance of each electrode was checked before recording and was kept below 40 *k*Ω. Additionally, electrode impedance levels were checked after every third block of 50 sentences (approx. every 30 minutes) and reduced if necessary.

### 2.6. Data Preprocessing & Feature Extraction

#### 2.6.1. Eye-Tracking Preprocessing & Feature Extraction

##### Eye-Tracking Preprocessing

The eye tracker computed eye position data and identified events such as saccades, fixations, and blinks. Saccade onsets were detected using the eye-tracking software default settings: acceleration larger than 8000°/s2, a velocity above 30°/s, and a deflection above 0.1°. The eyetracking data consists of (*x, y*) gaze location entries for each individual time point (Figure 3b). Coordinates were given in pixels with respect to the monitor coordinates (the upper left corner of the screen was (0, 0) and down/right was positive). We provide this raw data as well as various engineered eyetracking features.

##### Eye-Tracking Feature Extraction

For this feature extraction, only fixations within the boundaries of each displayed word were extracted. A Gaussian mixture model was trained on the (y-axis) gaze data for each sentence to improve the allocation of eye fixations to the corresponding text lines. The number of text lines determined the number of Gaussians to be fitted within the model. Subsequently, each gaze data point was clustered to the matching Gaussian and the data were realigned. As a result, each gaze data point is clearly assigned to a specific text line. Data points distinctly not associated with reading (minimum distance of 50 pixels to the text) were excluded. Additionally, fixations shorter than 100 ms were excluded from the analyses, because these are unlikely to reflect fixations relevant for reading (Sereno and Rayner, 2003). On the basis of the GECO and ZuCo 1.0 corpora, we extracted the following features: (i) *gaze duration* (GD), the sum of all fixations on the current word in the first-pass reading before the eye moves out of the word; (ii) *total reading time* (TRT), the sum of all fixation durations on the current word, including regressions; (iii) *first fixation duration* (FFD), the duration of the first fixation on the prevailing word; (iv) *single fixation duration* (SFD), the duration of the first and only fixation on the current word; and (v) *go-past time* (GPT), the sum of all fixations prior to progressing to the right of the current word, including regressions to previous words that originated from the current word. See Figure 2 for a visualization of the feature ranges of each reading task. For each of these eye-tracking features, we additionally computed the pupil size. Furthermore, we extracted the number of fixations and mean pupil size for each word and sentence. Additionally, on sentence level, we extracted the mean and maximum saccade velocity, saccade amplitude and saccade duration. On word level, saccade velocity, amplitude, and duration were extracted for in-going, outgoing, as well as saccades within a word. Finally, on the sentence level, omission rate is calculated, representing the proportion of words which were not fixated within each sentence.

**Figure 2:**
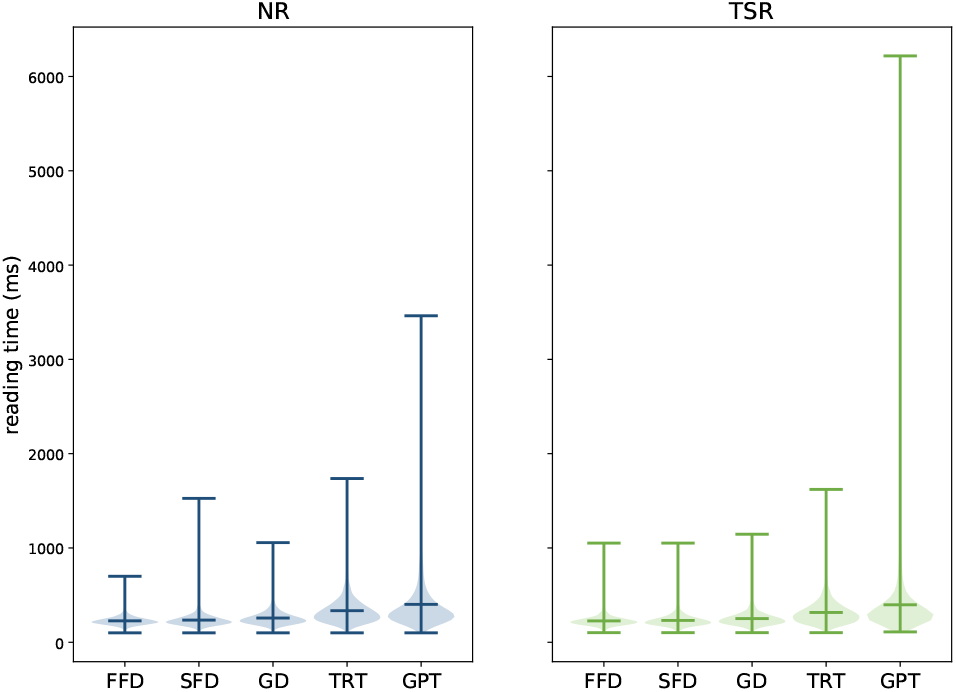
Violin plots showing means, distributions, and ranges of the reading time measures per word for each task and each eye-tracking feature (x-axis) in milliseconds.

#### 2.6.2. EEG Preprocessing & Feature Extraction

##### EEG Preprocessing

Before the EEG preprocessing, data from all 14 blocks (7 NR and 7 TSR) were first merged to avoid high predictive power based on the differences resulting from the preprocessing itself. To avoid loss of data by the subsequent automated preprocessing pipeline, the files of each recording blocked were screened to exclude highly artifactual data. Therefore, each block was temporarily filtered using a 2Hz high-pass filter. Subsequently, outlying data points were removed if they exceeded a threshold of three standard deviations above or below the mean of the data. Only if the standard deviation of this temporarily pre-cleaned data was below a cut-off of 100 microvolt, the original corresponding block was used in the merging process. Applying this criterion, 4.02% of all blocks were excluded. The EEG preprocessing was conducted with the open-source MATLAB tool-box preprocessing pipeline Automagic (Pedroni et al., 2019), which combines state-of-the-art EEG preprocessing tools into a standardized and automated pipeline. The EEG preprocessing consisted of the following steps: First, bad channels were detected by the algorithms implemented in the EEGlab plugin `clean_rawdata`^4^. A channel was defined as a bad electrode when recorded data from that electrode was correlated at less than 0.85 to an estimate based on other channels. Furthermore, a channel was defined as bad if it had more line noise relative to its signal than all other channels (4 standard deviations). Finally, if a channel had a longer flat-line than 5 seconds, it was considered bad. These bad channels were automatically removed and later interpolated using a spherical spline interpolation (EEGLAB function eeg_interp.m). The interpolation was performed as a final step before the automatic quality assessment of the EEG files. Next, data were filtered using a 2 Hz high-pass filter and line noise artifacts were removed by applying Zapline (de Cheveigné, 2020), removing seven power line components. Subsequently, independent component analysis (ICA) was performed. Components reflecting artifactual activity were classified by the pre-trained classifier ICLabel (Pion-Tonachini et al., 2019). Components that were classified as any class of artifacts (line noise, channel noise, muscle activity, eye activity, and cardiac artifacts) with a probability higher than 0.8 were removed from the data. Subsequently, residual bad channels were excluded if their standard deviation exceeded a threshold of 25*μV*. Very high transient artifacts (> 100*μV*) were excluded from calculating the standard deviation of each channel. However, if this resulted in a significant loss of channel data (> 50%), the channel was removed from the data. After this, the pipeline automatically assessed the quality of the resulting EEG files based on four criteria: First, a data file was marked as bad-quality EEG and not included in the analysis if the proportion of high-amplitude data points in the signals (> 30*μV*) was larger than 0.20. Second, more than 20% of time points showed a variance larger than 15*μV* across channels. Third, 30% of the channels showed high variance (> 15*μV*). Fourth, the ratio of bad channels was higher than 0.3. After Automagic preprocessing, 13 electrodes in the outermost circumference (chin and neck) were excluded from further processing as they capture little brain activity and mainly record muscular activity. The discarded electrode labels were E1, E8, E14, E17, E21, E25, E32, E48, E49, E56, E63, E68, E73, E81, E88, E94, E99, E107, E113, E119, E125, E126, E127, and E128. Additionally, 10 EOG electrodes were separated from the data and not used for further analysis, yielding a total number of 105 EEG electrodes. Subsequently, the data was converted to a common average reference.

##### EEG & Eye-Tracking Synchronization

In a next step, the EEG and eyetracking data were synchronized using the “EYE-EEG” toolbox (Dimigen et al., 2011) to enable EEG analyses time-locked to the onsets of fixations and saccades, and subsequently segment the EEG data based on the eye-tracking measures. The synchronization algorithm first identified the “shared” events. Next, a linear function was fitted to the shared event latencies to refine the start- and end-event latency estimation in the eye tracker recording. Finally, the synchronization quality was ensured by comparing the trigger latencies recorded in the EEG and eye-tracker data. All synchronization errors did not exceed 2 ms (i.e., one data point). Remaining eye artifacts in data were removed with Unfold toolbox (Ehinger and Dimigen, 2019) according to a method described in (Pfeiffer et al., 2020) The effect of this preprocessing can be seen in Figure 3d.

**Figure 3:**
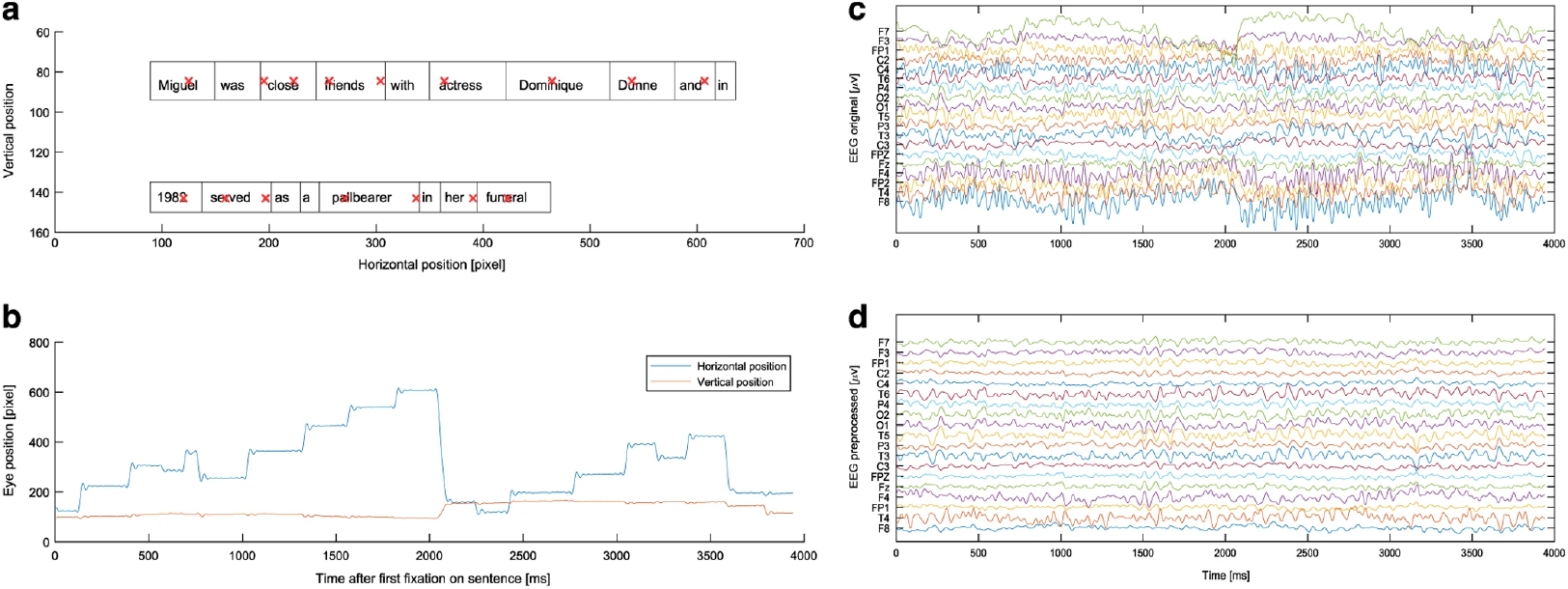
Visualization of eye-tracking and EEG data for a single sentence. (a) Prototypical sentence fixation data. Red crosses indicate fixations; boxes around the words indicate the wordbounds. (b) Fixation data plotted over time. (c) Raw EEG data during a single sentence. (d) Same data as in (c) after preprocessing.

##### EEG Feature Extraction

To compute oscillatory power measures, we bandpass filtered the continuous EEG signals across an entire reading task for four different frequency bands, resulting in a time-series for each frequency band. The distinct frequency bands were determined as follows: *theta_1* (4-6 Hz), *theta_2* (6.5-8 Hz), *alpha_1* (8.5-10 Hz), *alpha_2* (10.5-13 Hz), *beta_1* (13.5-18 Hz), *beta 2* (18.5-30 Hz), *gamma_1* (30.5-40 Hz), *gamma_2* (40.5-49.5 Hz). Afterwards, we applied a Hilbert transformation to each of these time-series resulting in a complex time series. The Hilbert phase and amplitude estimation method yields results equivalent to sliding window Fourier transformation and wavelet approaches (Bruns, 2004). We chose specifically the Hilbert transformation to maintain temporal information for the amplitude of the frequency bands to enable the power computation of the different frequencies for time segments defined through fixations in the eye-tracking data. Finally, for each sentence as well as for each word within each sentence, and for each frequency band, the EEG features consist of a vector of 105 dimensions (one value for each EEG channel). On the level of individual words, these frequency band power features were calculated based on fixations of GD, TRT, FFD, SFD, and GPT (see above). For each EEG feature, all channels were subject to an artifact rejection criterion of 90*μV* to exclude trials with transient noise. To descriptively compare the EEG activity and the extracted frequency band power between the NR and TSR sentences, the differences (NR minus TSR) for the different sentence-level EEG features are plotted in Figure 4.

**Figure 4:**
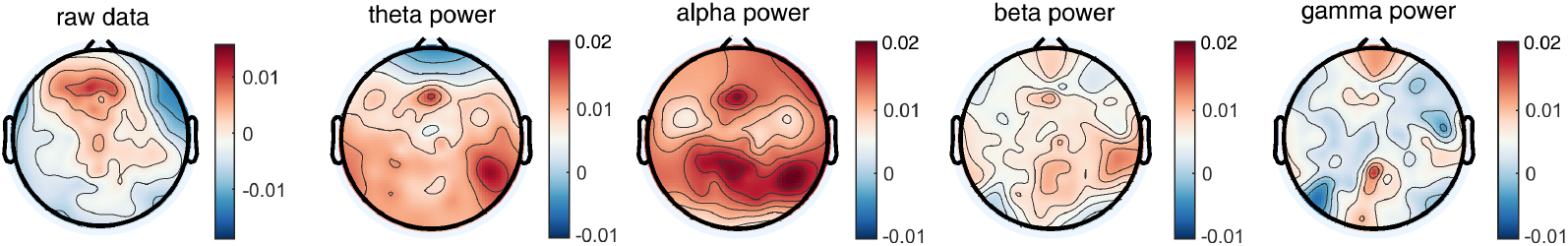
Topography plots showing the mean EEG activity across all subjects from ZuCo 2.0 of the difference between the tasks (NR minus TSR) (scalp viewed from above, nose at the top).

### 2.7. Data Access

The raw and preprocessed EEG and eye-tracking data, as well as the features extracted from the preprocessed EEG and eye-tracking are provided for this benchmark. For the training data, the information about the task (normal reading or task-specific reading) is also available. Please note that for the held-out test dataset, we can only provide the preprocessed data and the extracted features. As the raw data were collected in different blocks of normal reading and task-specific reading, the participants could otherwise infer the outcome from the block separation. All the data can be accessed via OSF: https://osf.io/d7frw/.

## 3. Benchmark Task

### 3.1. Task Definition

We propose an ML benchmark for reading task identification. As described in Section 2, the ZuCo corpus provides data from two reading paradigms, normal reading (NR) and task-specific annotation reading (TSR). Consequently, we frame the problem as binary classification task with labels *Y* ∈ {NR, TSR}. The training data consists of sentences labeled depending on which reading task they belonged to during the experiment. Each sentence is represented by a feature set *X*. The input features should be eye-tracking or EEG features, or a combination thereof.

The goal of the benchmark task is to build a binary classifier *h* to predict the label *Y* for each sentence given only the features *X*:

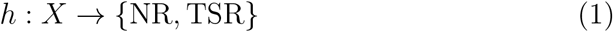

Due to the naturalistic experiment design and the co-registration of EEG and eye movement signals, feature extraction is possible on various levels. There are no restrictions to the type and dimension of the input features or the model.

### 3.2. Performance Metrics

The classifier’s performance is evaluated by the classification accuracy, defined as the number of correct predictions divided by the total number of predictions. Since previous results have shown high performance on models trained and tested within-subject but low performance on cross-subject models (Hollenstein et al., 2021c), this benchmark aims to address this gap by focusing on the latter to improve the inter-subject generalization capabilities of the models. We propose a cross-subject evaluation, where each subject in the held-out testset is evaluated by a model trained on all subjects in the training split (i.e., the original ZuCo 2.0 dataset). Therefore, the main benchmark metric is defined as the mean classification accuracy across all subjects in the testset. As a second metric, we choose the F1-measure. In our classification setup, we do not distinguish between a positive and a negative class, i.e., there is no clear majority or minority class. For that reason, we choose to evaluate our classifier using the macro-averaged F1-scores. The benchmark task is evaluated on models from the following three categories: models trained on EEG features, models trained on eye-tracking features, and models trained on a combination of EEG and eye-tracking features.

### 3.3. Benchmark Setup

We host the ZuCo benchmark on Eval-AI (Yadav et al., 2019) – an open source AI challenge platform for evaluating and comparing machine learning and artificial intelligence algorithms. The reading task classification challenge page is available here: https://eval.ai/web/challenges/challenge-page/1444/overview. This solution will help other researchers to participate in our machine learning challenge and enable us to automate the evaluation of the future submissions.

#### Evaluation Strategy

Researchers that want to participant in the benchmark task can submit predictions from their models for the hidden testset. We specified the challenge configuration, evaluation code, and information about the data splits. Predictions for the testset labels can be submitted in the JSON file.

#### Leaderboard

The public leaderboard will include the scores on the chosen evaluation metrics as well as references to upcoming publications. Upon submission, the predictions will be handed over to challenge-specific workers that compare the predictions against corresponding ground-truth labels using the custom evaluation script provided by our team.

## 4. Baseline Methods

### 4.1. Textual Baselines

We set three minimal baselines for this benchmark task: (i) a random baseline, (ii) a word embedding baseline, and (iii) a text difficulty baseline.

#### Random Baseline

We compute a random baseline to assess the chance level of predicting the correct class. We randomly sample the labels according to the distribution of the training data. That means the label NR is chosen with a probability of 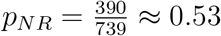 and TSR is chosen with *p_TSR_* = 1 – *p_NR_* ≈ 0.47.

#### Word Embedding Baseline

We compare our models to a textual baseline as a sanity check to ensure the sentences in the data are not easily separable merely by their linguistic characteristics. For this purpose, we use pre-trained textual representations, namely the state-of-the-art contextualized BERT word embeddings (Devlin et al., 2019). We concatenate the embeddings of all words in a sentence feed the into the LSTM model.

#### Text Difficulty Baseline

We also provide a baseline based on text readability. Although the sentences for both reading tasks were chosen to be of similar length and from the same text genre, we want to ensure that both tasks are not separable merely by the difficulty of the sentences. Therefore, we implement a text difficulty baseline, which classifies the sentences into NR and TSR based on their Flesch reading ease score (FRE; Flesch, 1948). This score indicates how difficult an English text passage is to understand based on the average number of words in a sentence and the average number of syllables in a word:

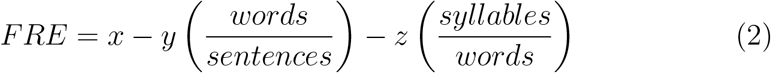

where *x, y* and *z* are language-specific weighting factors (for English *x* = 206.835, *y* = 1.015, *z* = 84.6). We compute FRE scores for each of the English sentences in the ZuCo data. Figure 5 shows the distribution of the FRE across the sentences of ZuCo 2.0.

**Figure 5:**
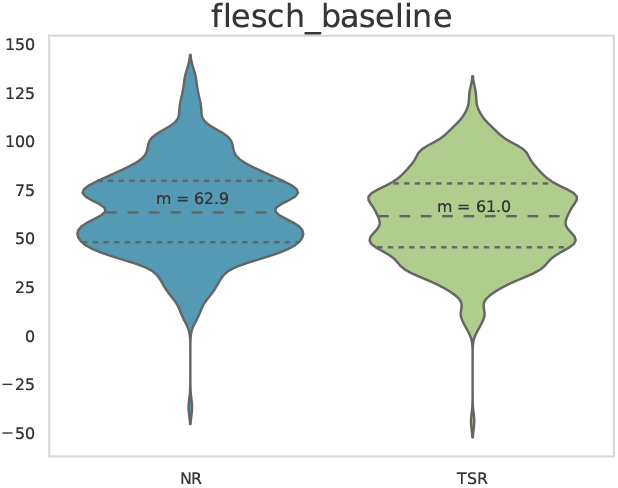
Flesch reading ease (FRE) scores for the NR and TSR sentences used in the ZuCo 2.0. dataset

### 4.2. EEG & Eye-tracking Models

We also present a set of initial models using EEG and eye-tracking features as a starting point for future models^5^. For each sentence in the dataset, the model input is composed of a vector of eye-tracking and/or EEG features corresponding to a single sentence in the dataset. Each sample in the training set is labelled with the reading task it was recorded in, normal reading (NR) or task-specific reading (TSR). We investigate the potential of using sentence-level eye-tracking and EEG features for the reading task classification. Hollenstein et al. (2021c) compared sentence-level and word-level features for this task previously and showed that sentence-level features perform better. However, challenge participants are also invited to use wordlevel and other features (see discussion in Section 6 for suggestions). The advantages of sentence-level features consist of the possibility of using simpler machine learning models and reduced training times (Hollenstein et al., 2021c). Sentence-level features are defined as metrics aggregated over all words in a given sentence.

#### Eye-tracking Features

We include two types of sentence-level eye-tracking features. The features are summarized in Table 5. First, the fixation-based features - omission rate, number of fixations and reading speed - are aggregated metrics normalized by sentence length, i.e., the number of words in a sentence. Analogous to the word-level models, we also include saccade-based features. These include the mean and maximum duration, velocity and amplitude across all saccades that occurred within the reading time of a give sentence. We test these features individually and combined to investigate the performance increase achieved by adding more features.

**Table 5:** Sentence-level eye-tracking features. We use the combination of all features for our models.

| Name | Definition | Values |
| --- | --- | --- |
| FIXATION FEATURES |  |  |
| omission_rate | Percentage of words in a sentence that is *not* fixated | 1 |
| fixation_number | Number of fixations in the sentence divided by the number of words | 1 |
| reading_speed | Sum of the duration of all fixations in the sentence divided by the number of words | 1 |
| SACCADE FEATURES |  |  |
| mean_sacc_dur | Sum of the duration of all saccades in the sentence divided by the number of words | 1 |
| max_sacc_dur | Maximum saccade duration per sentence | 1 |
| mean_sacc_velocity | Sum of the velocity of all saccades in the sentence divided by the number of saccades | 1 |
| max_sacc_velocity | Maximum saccade velocity per sentence | 1 |
| mean_sacc_amplitude | Sum of the amplitude of all saccades in the sentence divided by the number of saccades | 1 |
| max_sacc_amplitude | Maximum saccade amplitude per sentence | 1 |
| COMBINED FEATURES |  |  |
| Combined ET features | Concatenation of all eye-tracking features | 9 |

#### EEG Features

The sentence-level EEG features take into account the EEG activity over the whole sentence duration (even when no words were fixated). We aggregate over the preprocessed EEG signals of the full reading duration of a sentence. Each subfrequency band (e.g., *alpha_1* and *alpha_2*) were averaged to get one power measure for each frequency band, i.e., *theta* (4-8 Hz), *alpha* (8.5-13 Hz), *beta* (13.5-30 Hz), and *gamma* (30.5-49.5 Hz). The sentence-level EEG features are described in Table 6. We experiment with both aggregate metrics, i.e., the mean across all electrodes, and individual electrode features.

**Table 6:** Sentence-level EEG features.

| Name | Definition | Values |
| --- | --- | --- |
| MEAN FEATURES |  |  |
| theta_mean | Mean theta band features averaged over all electrodes | 1 |
| alpha_mean | Mean alpha band features averaged over all electrodes | 1 |
| beta_mean | Mean beta band features averaged over all electrodes | 1 |
| gamma_mean | Mean gamma band features averaged over all electrodes | 1 |
| eeg_means | Mean frequency band features averaged over all electrodes, resulting in 1 feature value for each of the 8 frequency bands | 8 |
| ELECTRODE FEATURES |  |  |
| electrode_features_theta | Mean theta1 and theta2 values of all 105 electrodes | 105 |
| electrode_features_alpha | Mean alpha_1 and alpha_1 values of all 105 electrodes | 105 |
| electrode_features_beta | Mean beta_1 and beta_1 values of all 105 electrodes | 105 |
| electrode_features_gamma | Mean gamma_1 and gamma_1 values of all 105 electrodes | 105 |
| electrode_features_all | Concatenation of the four features above | 420 |
| COMBINED FEATURES |  |  |
| ET & EEG mean features | Concatenation of sent_gaze_sacc and eeg_means | 17 |

Examples of these features across all subjects, split by class (normal reading vs. task-specific reading) are shown in Figure 6 for ZuCo 2.0.

**Figure 6:**
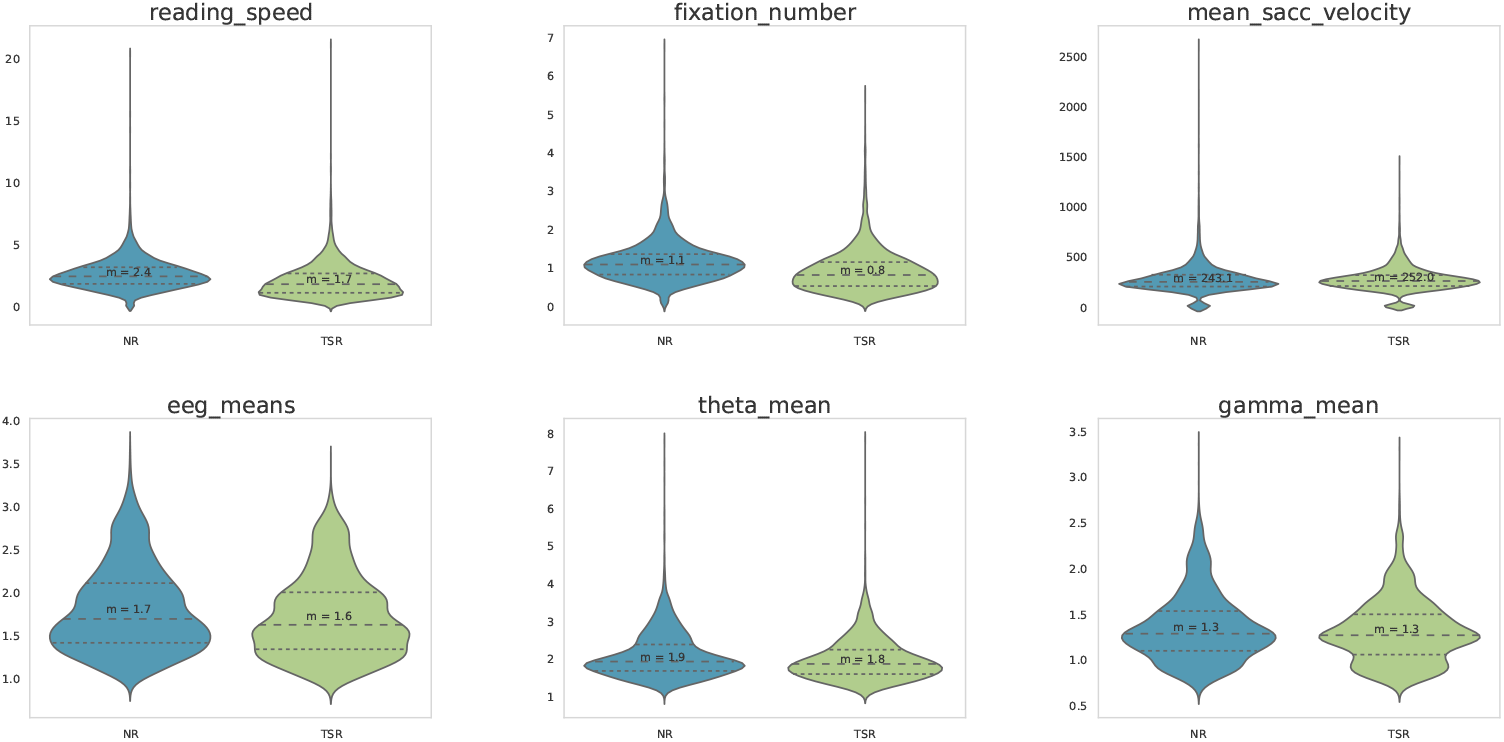
Examples of feature distributions across all subjects for the NR and TSR sentences included in the ZuCo 2.0. dataset

#### Principal Component Analysis

We additionally employed principal component analysis (PCA) to reduce the high dimensionality of the EEG features. In an initial attempt, we fitted PCA on all training subjects and applied it to both the training and test split. However, no significant improvements in the classification accuracy could be observed. Thus, we fitted PCA to each subject individually. In order to determine the number of components of PCA, which will later be used by the classifier, we only consider the subjects in the training data to prevent overfitting to the test subjects. Thereby, we fit PCA for each of those subjects separately and calculated the number of components that explain 95% of the variance. We then choose the number of components of PCA as the median over all subjects in the training data, which makes it robust against outlier subjects. With this way, PCA reduced the dimensionality of training and test data from 105 to 41. Figure 7 shows that the amount of variance explained by the first components varies greatly between subjects. For instance, the first component accounts for approx. 24% of the variance for subject YTL, whereas it accounts for 49% of the variance for subject YAC.

**Figure 7:**
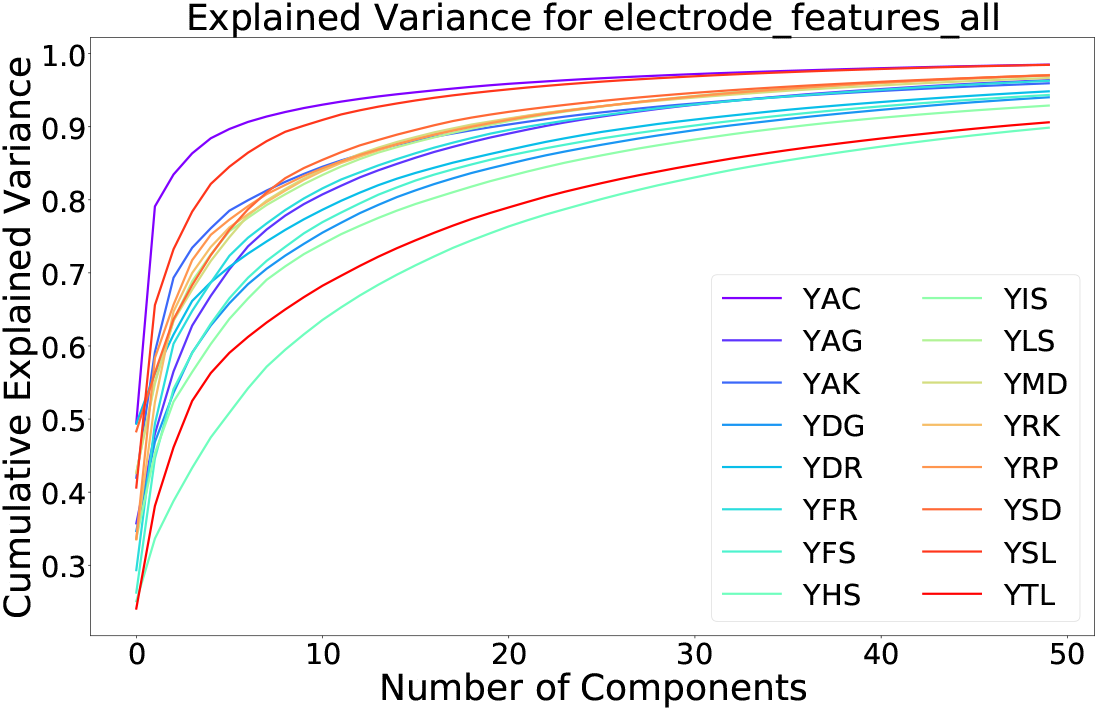
Variance explained with increasing number of PCA components for the training subjects in ZuCo 2.0.

To analyze how much the original features, i.e., the individual electrodes, influence the principal components, we fit a PCA for each subject of the training data, such that the resulting components explain 95% of the variance. Assuming we have *n* original features and *m* principal components c, where each component is a linear combination of the original features, i.e., 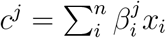, *j* ∈ 1… *m*. We then extract the amount of variance explained (*v^j^*) by each component *c^j^* and its weights 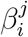. We sum up all 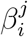 weighted by *v^j^*, such that the resulting 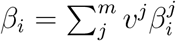 represents the relevance of feature *x_i_*.

Following this procedure, we split the results into frequency bands and present the corresponding topography plots averaged over all training subjects in ZuCo 2.0 in Figure 8.

**Figure 8:**
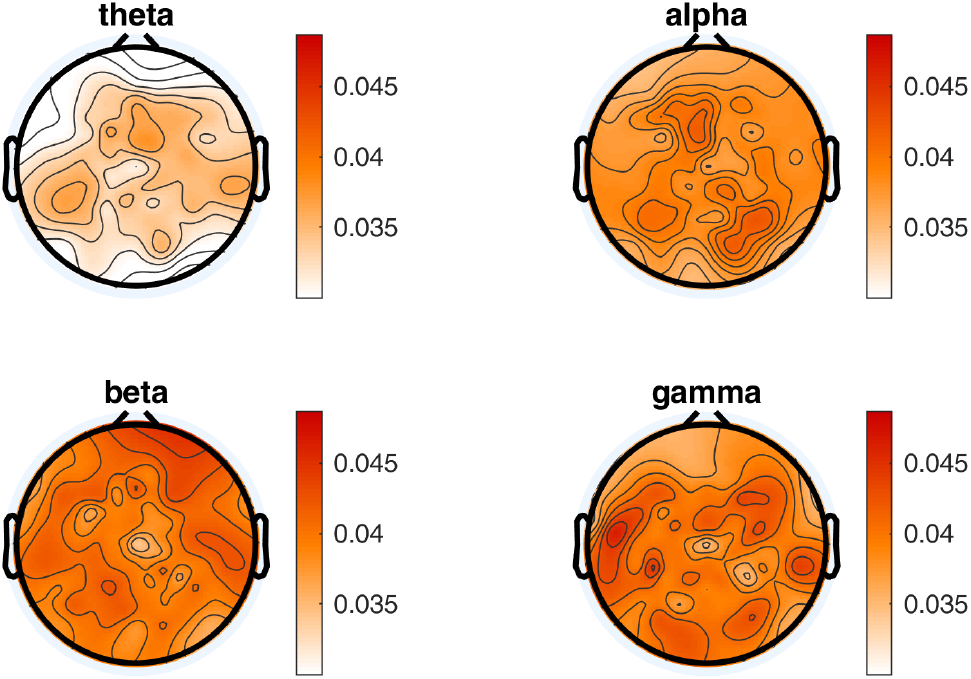
Topographical distribution of electrode importance for the principle components, divided into the 4 different frequency bands. Electrode importance is calculated by determining the influence of each electrode on the principle components and weighting them by amount of explained variance.

#### Model

The input to the sentence-level model is a single vector representing each sentence. We scale the feature values to a range between {0, 1}. We train a support vector machine for classification with a linear kernel. We use the `scikit-learn` SVC implementation^6^. For the cross-subject evaluation, the models are trained on all samples from all subjects in ZuCo 2.0 and tested on the samples from new subjects in the held-out testset.

## 5. Results

### 5.1. Results of Textual Baselines

As described in the previous section, we set three minimal baselines for this benchmark task: (i) a random baseline, i.e., chance level for binary classification, (ii) a word embedding baseline, namely BERT word embeddings, and (iii) a text difficulty baseline, based on the Flesch reading ease score (FRE). The random baseline for binary classification is at 0.50 accuracy. The word embedding baseline yield a classification accuracy of 0.65 for ZuCo 2.0. The text difficulty baseline is also above random performance with a classification accuracy of 0.53 for ZuCo 2.0. Table 7 shows the accuracy and F1-score for all baselines.

**Table 7:** The mean accuracy and F1-score over all subjects for each feature-set in the benchmark task.

| Feature Set | Accuracy | F1 |
| --- | --- | --- |
| Random | 0.50 | 0.50 |
| FRE Baseline | 0.53 | 0.35 |
| BERT Baseline | 0.65 | 0.64 |
| Eye-tracking features | 0.69 | 0.67 |
| Eye-tracking and EEG mean features | 0.68 | 0.66 |
| Concatenated EEG electrode features | 0.55 | 0.46 |
| Concatenated EEG electrode features (with PCA) | 0.58 | 0.56 |

### 5.2. Results of EEG & Eye-tracking Models

As described in Section 3, we consider three different feature sets, EEG, eyetracking, and the combination of all features. For each feature set and each subject, we report the accuracy and the F1-score. For each subject in the hidden testset, we compute the results via bootstrapping, sampling 500 times with replacement, and using a sample size equal to the original data. For all results, we report the comparison to the random and textual baselines as well as the 95% confidence intervals for each subject. Table 7 shows a summary of the results. The corresponding tables with the detailed numbers for all subjects and feature sets are shown in the Appendix A.

First, the results for the eye-tracking features are shown in Figure 9. These results clearly show all subjects outperforming the random baseline and FRE baseline except for one subject each for accuracy and F1-score. All subjects except one perform better than the random baseline, and three subjects perform significantly better than the BERT word embedding baseline. The mean accuracy across all subjects in the testset is 0.69, and the mean F1-score is 0.67. Furthermore, the results for the combined eye-tracking and EEG mean feature set in Figure 10 do not yield an increase in performance compared to using only the eye-tracking features (mean accuracy: 0.68; F1-score: 0.66). Interestingly, the best and worst performing subjects vary between different feature combinations, and between accuracy and F1-score.

**Figure 9:**
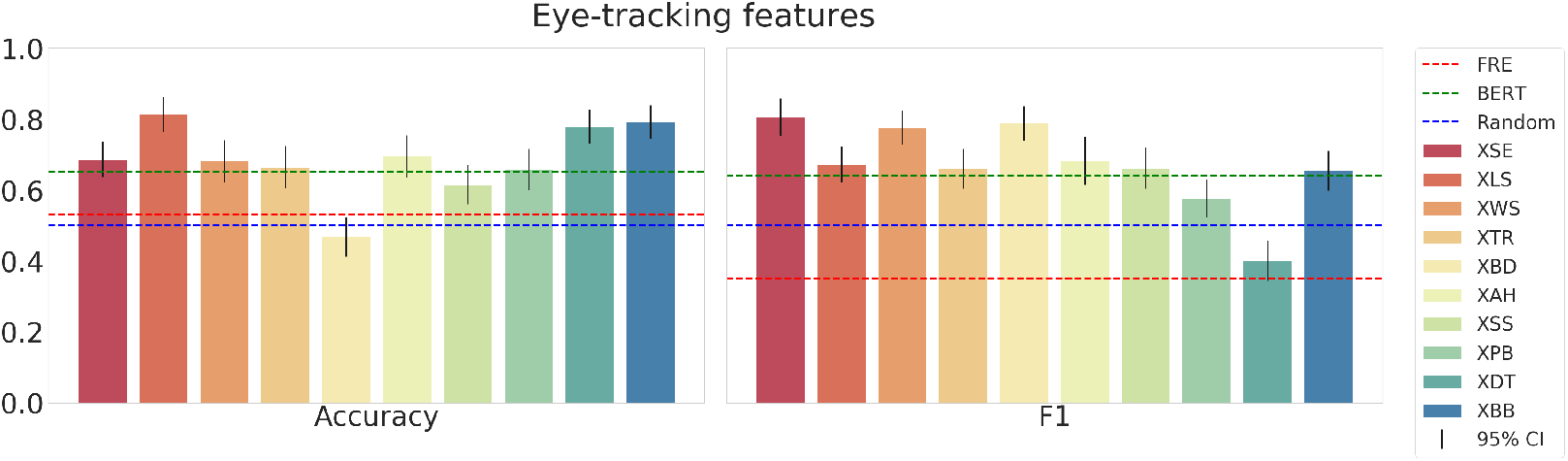
The mean accuracy (left), mean F1-score (right) with corresponding 95% confidence intervals and textual baselines are plotted for each subject in the held-out test dataset using the concatenated eye-tracking features.

**Figure 10:**
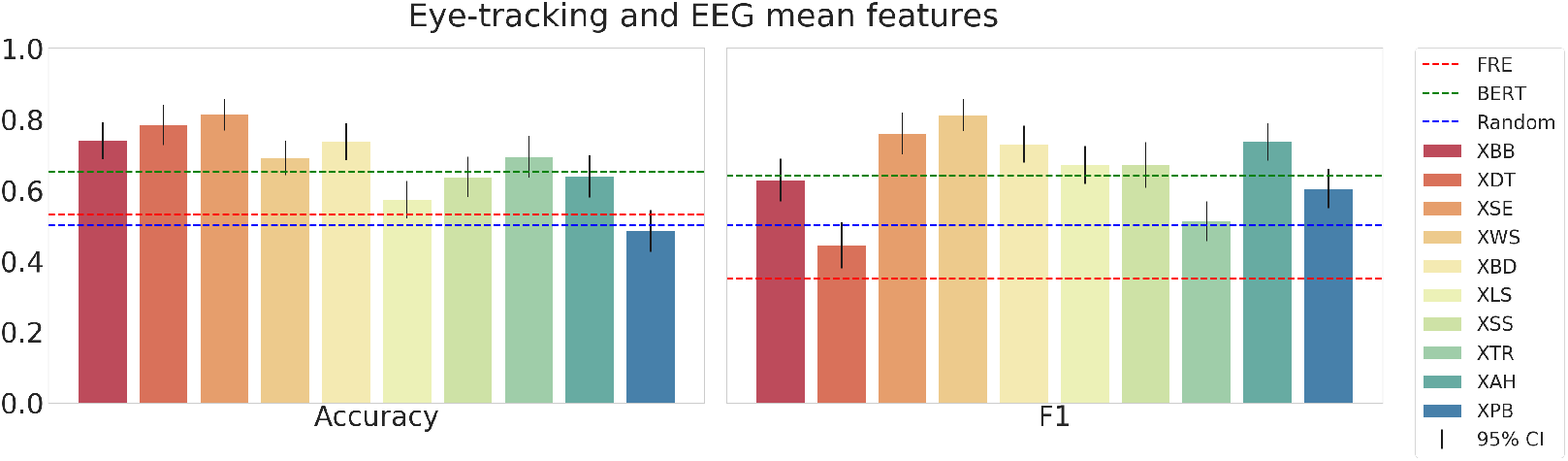
The mean accuracy (left), mean F1-score (right) with corresponding 95% confidence intervals and textual baselines are plotted for each subject in the held-out test dataset using the eye-tracking and EEG mean features.

Next, we show the results using the concatenated EEG electrode features in Figure 11. With this feature set, the mean accuracy across all subjects in the testset is 0.55, and the mean F1-score is 0.46. The accuracy scores are notably higher than for the F1-score. Finally, when using the same features but applying the PCA preprocessing, the models yield the results presented in Figure 12. The scores for the accuracy are similar but have a slightly higher mean of 0.58 (compared to 0.55 without PCA). However, the F1-scores with PCA are significantly higher with a mean of 0.56 (compared to 0.46 without PCA). While with these EEG electrode features the models outperform the random and text difficulty baseline for some test subjects, they do not achieve to outperform the strong embedding baseline.

**Figure 11:**
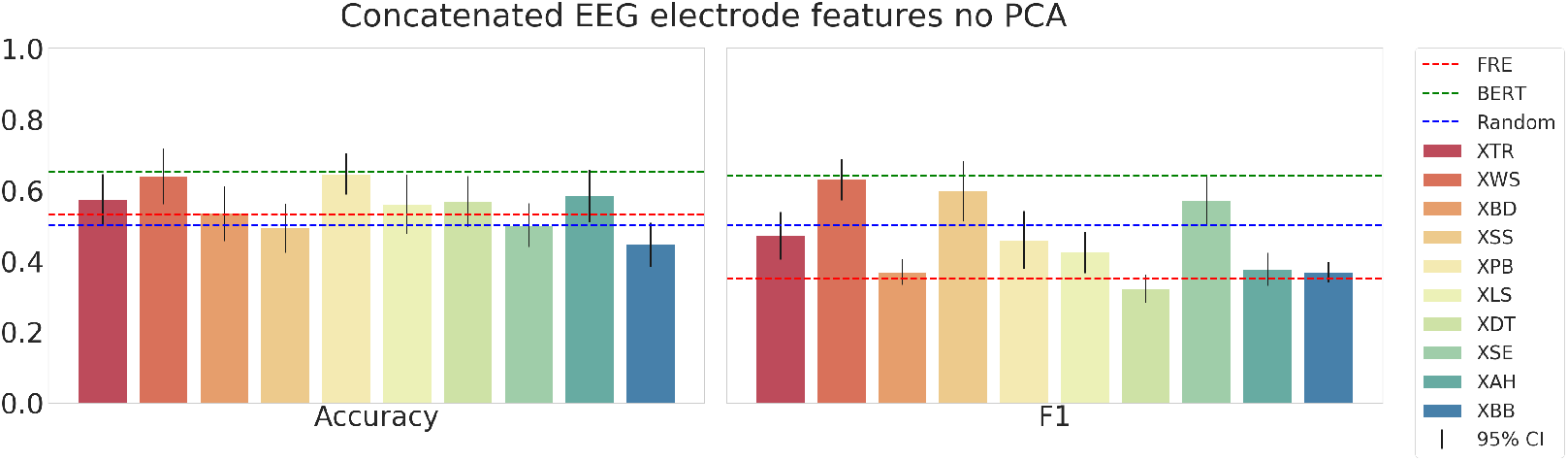
The mean accuracy (left), mean F1-score (right) with corresponding 95% confidence intervals and textual baselines are plotted for each subject in the held-out test dataset using the concatenated EEG electrode features without PCA pre-processing.

**Figure 12:**
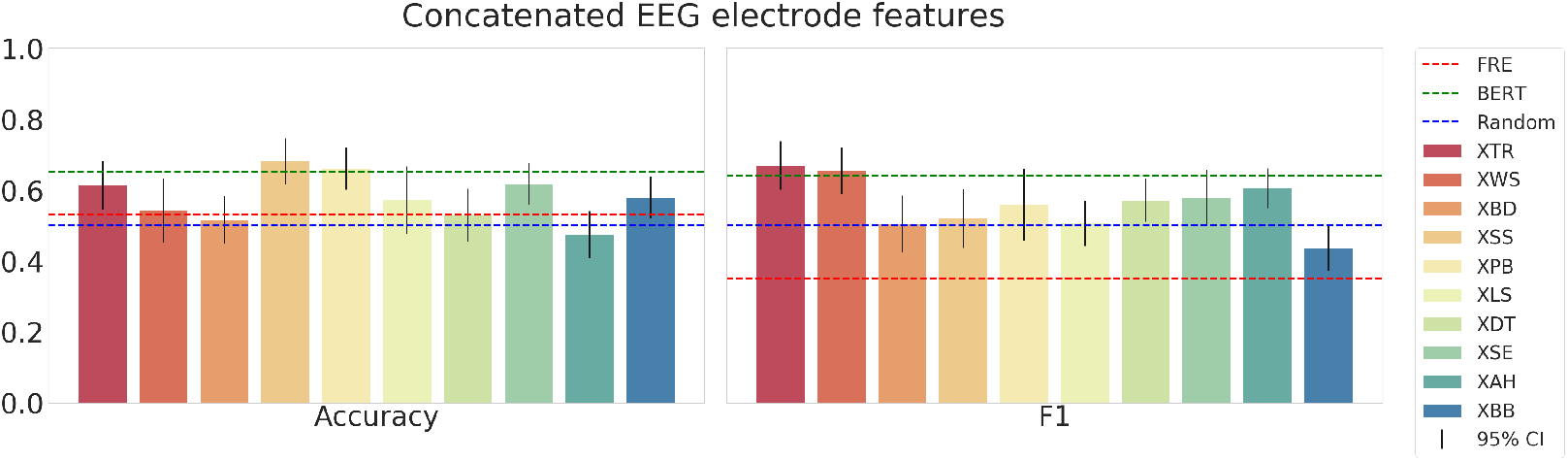
The mean accuracy (left) mean, F1-score (right) with corresponding 95% confidence intervals and textual baselines are plotted for each subject in the held-out test dataset using the concatenated EEG electrode features after pre-processing with PCA.

## 6. Discussion

The present benchmark challenge has the main goal of advancing reading task classification through eye-tracking and EEG data. The challenge participants are invited to develop ML models to identify whether subjects are reading a sentence with the goal of reading comprehension (i.e., normal reading) or whether the subjects are reading a sentence to search for a specific semantic relation in the sentence (i.e., task-specific reading). The objective is to investigate which eye movement and brain activity features are most suited to solve this problem. Understanding the physiological aspects of the reading process (i.e., the cognitive load and reading intent) can advance our understanding of human language processing. On the other hand, natural language processing and machine learning would benefit, as classifiers that outperform current textual baselines could improve the quality and process of collecting annotated data (e.g., through gaze-aided unsupervised labelling).

Several previous studies have used ML models to accurately perform a reading task classification. Cole et al. (2011) used eye-tracking data to discriminate between a scanning task and a reading comprehension task. Furthermore, Biedert et al. (2012) developed a real-time classifier able to distinguish reading from skimming patterns. In a related study, Kelton et al. (2019) investigated the influence of different content and tasks on the performance to determine whether subjects are reading or skimming a news article. Other neuroimaging methods such as fMRI have been combined with eye-tracking to examine the neural basis of sentence comprehension (e.g., Bonhage et al., 2015) or the discrimination between normal and non-word text (Choi et al., 2014). In another fMRI study, which simultaneously recorded eye-tracking data, Ceh et al. (2021) observed that internally and externally directed cognition are characterized by distinct brain activity. In addition, several research groups provide publicly available fMRI data to study naturalistic reading comprehension (Pereira et al., 2018; Dehghani et al., 2017; Lopopolo et al., 2018; Shain et al., 2020; Nastase et al., 2021). While functional MRI has a better spatial resolution compared to EEG, is a very costly method with restricted real-life usability. Whereas eye-tracking and EEG systems are of lower cost and can be used in more naturalistic situations. Several other publicly datasets recorded eye-tracking (e.g., Cop et al., 2017; Luke and Christianson, 2017; Jäger et al., 2021) or EEG from continuous speech stimuli (e.g., Broderick et al., 2018; Brennan and Hale, 2019). These datasets provide the possibility to improve and evaluate machine learning systems for NLP. However, to the best of our knowledge, the ZuCo dataset is the only publicly available dataset that features simultaneous eye movement and EEG data recorded in a naturalistic reading setup. Thus, ZuCo is specifically tailored to leverage EEG and eye-tracking data to improve natural language processing tasks in a naturalistic setting. The field of machine learning contains a range of tasks on different modalities such as language (text), computer vision (video, images), and speech recognition (audio). Recently, Akbari et al. (2021) have shown superior performance of ML models with multimodal representations on downstream tasks such as image classification. Therefore, from an NLP perspective, another extension to this benchmark could be to investigate whether leveraging multimodal embeddings is beneficial for reading task classification.

In a recent study, the ZuCo data has been used already for reading task identification (Mathur et al., 2021) using a complex convolutional network, which is evaluated on a fixed cross-subject scenario on the sentences from ZuCo 2.0. However, the relatively poor performance of their model evaluated in a fixed cross-subject scenario, still leaves room for improvement and opens research questions regarding the selection of features. Hollenstein et al. (2021c) have recently presented extensive work on reading task classification, corroborating the advantages of the ZuCo dataset for this ML task. The authors found that, while high accuracy can be achieved on within-subject models, the performance drops for cross-subject evaluations. A current bottleneck of machine learning is the lack of generalization capabilities of these models, meaning that the models perform poorly on data from other domains that are not included in their training data. For instance, ML models perform less accurately across languages, across image or text domains, or across subjects. The latter is of great importance in neuroscientific research which aims at a principled understanding of human brain activity as a response to complex stimuli (Nastase et al., 2019), as well as for practical applications such as brain-computer interfaces (Chiang et al., 2019). Specifically, when trained on physiological data, the rules identified by ML models for a given task ideally hold for the entire population. Considering the ever-increasing complexity of ML models due to their large number of parameters, they are prone to overfit to their training set (which does not characterize the entire population), leading to spurious correlations. Therefore, to validate the gained insights on the physiological data, ML models need to be evaluated on held-out subjects as a proxy to the model’s generalization capability. These results inspired the proposed benchmark based on the ZuCo dataset. The benchmark task and baseline models follow the rules suggested by Scheinost et al. (2019) to take into account subject-specific differences in predictive modeling.

In the current paper, we provide evidence that both eye-tracking and brain activity data can improve reading task classification compared to purely text-based baselines. In the current work, the best-performing model is based on sentence-level eye-tracking features. Combining eye-tracking and EEG mean features yields promising results, but not better than eye-tracking only. Therefore, there are possible gains in performance to be achieved by more sophisticated combinations of eye movement and brain activity features. There are various ways to leverage eye-tracking and EEG data. Currently, we extracted high-level eye-tracking features based on fixations (e.g, number of fixations and omission rate) and on saccades (e.g., mean velocity and maximum amplitude). The ZuCo dataset provides additional reading-related features such as mean fixation duration, total reading time or go-past time, but also pupil size information or even the raw data could be used in future approaches. Using raw data has shown great promise to model eye-tracking data (e.g., Jäger et al., 2020), and one of the main advantages of the ZuCo dataset is that it allows feature extraction on different levels. Moreover, our EEG features include mean features aggregated over all electrodes as well as electrode-based frequency measures, which have been shown to improve NLP tasks in the past (Hollenstein et al., 2019a; Sun et al., 2020; Hollenstein et al., 2021b; Wang and Ji, 2021). Nonetheless, we want to highlight that preprocessed EEG data permits the examination of additional measures, such as source-level based features (e.g., source-level power estimates) and functional connectivity measures at the level of the underlying neuronal generators. Other EEG analysis methods allow the extract measures of spatiotemporal dynamics of brain activity (e.g., microstates) (Michel and Koenig, 2018) and event-related potentials such as N400 components (Brouwer et al., 2017; Frank et al., 2013). Interestingly, Hollenstein et al. (2021c) found that gamma band features worked best in a within-subject setting. However, we found that concatenating all EEG electrode features is more beneficial in a cross-subject setting. Finally, the cross-subject performance can be further increased by using a dimensionality reduction (PCA) on the concatenated EEG features. Future methods could focus on new approaches for EEG feature selection and aggregation.

The simultaneous recording of EEG and eye-tracking allows us to investigate specific feature sets on different levels of analysis, e.g., sentence level, word level, fixation level. Additionally, the naturalistic setup of the experiments used in this work are crucial for this benchmark task and for neuroscience in general (Nastase et al., 2020). Not only does it increase the ecological validity of the recordings by allowing natural reading without controlling the individual reading speed, but it also supports the extraction of signals on various linguistic levels (Hasson and Egidi, 2015; Brennan, 2016; Hamilton and Huth, 2020; Kandylaki and Bornkessel-Schlesewsky, 2019; Alday, 2019). Frey et al. (2018) investigated how two different reading tasks modulate both eye movements and brain activity. In line with our findings, their results show that eye movement patterns were top-down modulated by different task demands. Moreover, their brain activity analysis suggests that the decision-making process during task-specific reading elicits a greater load in working memory than the one generated in a normal reading task. In summary, eye-tracking and EEG data offer an immensely diverse amount of potential measures, which might contain unique valuable information. Thus, we aim to inspire benchmark challenge participants to explore and extract alternative features from the available preprocessed data.

## 7. Conclusion

We presented a new ML benchmark using eye-tracking and EEG data to classify reading tasks. The goal of the benchmark challenge is to distinguish between normal reading and task-specific reading in a cross-subject evaluation scenario. We provide multiple initial models for this task and show that ML models trained on eye-tracking and EEG features can outperform strong textual baselines.

The standardized Zurich Cognitive Language Processing Corpus (ZuCo) dataset facilitates the creation of such a machine learning benchmark. We use the ZuCo 2.0 dataset as training data. To make our benchmark task more robust, we have additionally recorded further eye-tracking and EEG data from natural reading from additional subjects in a hidden testset. ZuCo’s rich structure and high-density coverage of simultaneous EEG and eye-tracking signals can also help to advance other areas that study the combination of gaze position and brain activity to identify variations in attention, reading patterns and reading intents, as well as participants’ compliance with the task demands and cross-subject variability.

Our dataset and benchmark setup allows us to easily add additional machine learning tasks to the leaderboard in the future. For instance, we can add additional NLP tasks since the ZuCo datasets provide ground truth labels for sentiment analysis or relation detection from text. Additionally, adding tasks such as eye movement and ERP prediction would be beneficial for various research communities. For example, the prediction of eye movement patterns has gained interest also in the NLP community (Hollenstein et al., 2021a). The main goal of this work is to create a platform for discussion and future research on a common benchmark task for reading task classification based on eye movement and brain activity data. We hope that this benchmark allows other researchers to make progress in this interdisciplinary research field.

## Acknowledgements

This work was partially supported by the Swiss National Science Foundation under grant 100014_175875 (Nicolas Langer) and by the German Federal Ministry of Education and Research under grant 01|S20043 (Lena A. Jäger).

## Appendix A. Appendix

**Table A.8:** Mean accuracy after bootstrapping the testset using 500 samples and a sample size equal to the original data for each subject in the held-out testset and each feature-set.

| Features | XBB | XDT | XLS | XPB | XSE | XTR | XWS | XAH | XBD | XSS |
| --- | --- | --- | --- | --- | --- | --- | --- | --- | --- | --- |
| sent_gaze_sacc | 0.66 | 0.47 | 0.69 | 0.62 | 0.81 | 0.66 | 0.78 | 0.7 | 0.79 | 0.68 |
| sent_gaze_sacc_eeg_means | 0.64 | 0.48 | 0.69 | 0.64 | 0.78 | 0.57 | 0.81 | 0.74 | 0.74 | 0.69 |
| electrode_features_all | 0.58 | 0.45 | 0.53 | 0.56 | 0.57 | 0.49 | 0.64 | 0.5 | 0.57 | 0.64 |
| electrode_features_all_pca | 0.47 | 0.58 | 0.52 | 0.57 | 0.61 | 0.68 | 0.66 | 0.62 | 0.53 | 0.54 |

**Table A.9:** Mean f1-score after bootstrapping the testset using 500 samples and a sample size equal to the original data for each subject in the held-out testset and each feature-set.

| Features | XBB | XDT | XLS | XPB | XSE | XTR | XWS | XAH | XBD | XSS |
| --- | --- | --- | --- | --- | --- | --- | --- | --- | --- | --- |
| sent_gaze_sacc | 0.65 | 0.4 | 0.67 | 0.58 | 0.81 | 0.66 | 0.78 | 0.68 | 0.79 | 0.66 |
| sent_gaze_sacc_eeg_means | 0.63 | 0.44 | 0.67 | 0.6 | 0.76 | 0.51 | 0.81 | 0.74 | 0.73 | 0.67 |
| electrode_features_all | 0.37 | 0.32 | 0.42 | 0.46 | 0.57 | 0.47 | 0.63 | 0.38 | 0.37 | 0.6 |
| electrode_features_all_pca | 0.44 | 0.57 | 0.51 | 0.56 | 0.58 | 0.67 | 0.65 | 0.61 | 0.5 | 0.52 |

**Table A.10:** Mean precision after bootstrapping the testset using 500 samples and a sample size equal to the original data for each subject in the held-out testset and each feature-set.

| Features | XBB | XDT | XLS | XPB | XSE | XTR | XWS | XAH | XBD | XSS |
| --- | --- | --- | --- | --- | --- | --- | --- | --- | --- | --- |
| sent_gaze_sacc | 0.7 | 0.48 | 0.7 | 0.74 | 0.81 | 0.69 | 0.82 | 0.71 | 0.83 | 0.71 |
| sent_gaze_sacc_eeg_means | 0.63 | 0.5 | 0.72 | 0.75 | 0.81 | 0.72 | 0.83 | 0.74 | 0.81 | 0.74 |
| electrode_features_all | 0.29 | 0.31 | 0.66 | 0.77 | 0.58 | 0.47 | 0.68 | 0.37 | 0.49 | 0.65 |
| electrode_features_all_pca | 0.44 | 0.57 | 0.52 | 0.58 | 0.59 | 0.66 | 0.68 | 0.61 | 0.5 | 0.52 |

**Table A.11:**
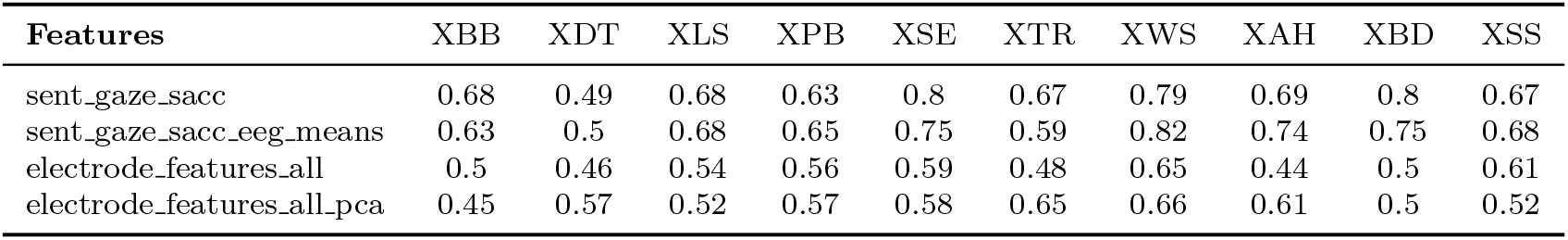
Mean recall after bootstrapping the testset using 500 samples and a sample size equal to the original data for each subject in the held-out testset and each feature-set.

| **Features** | XBB | XDT | XLS | XPB | XSE | XTR | XWS | XAH | XBD | XSS |
| --- | --- | --- | --- | --- | --- | --- | --- | --- | --- | --- |
| sent_gaze_sacc | 0.68 | 0.49 | 0.68 | 0.63 | 0.8 | 0.67 | 0.79 | 0.69 | 0.8 | 0.67 |
| sent_gaze_sacc_eeg_means | 0.63 | 0.5 | 0.68 | 0.65 | 0.75 | 0.59 | 0.82 | 0.74 | 0.75 | 0.68 |
| electrode_features_all | 0.5 | 0.46 | 0.54 | 0.56 | 0.59 | 0.48 | 0.65 | 0.44 | 0.5 | 0.61 |
| electrode_features_all_pca | 0.45 | 0.57 | 0.52 | 0.57 | 0.58 | 0.65 | 0.66 | 0.61 | 0.5 | 0.52 |

## Footnotes

1 Benchmark data available here: https://osf.io/d7frw/

2 Code for baseline methods available here: https://github.com/norahollenstein/zuco-benchmark

3 Data available here: https://osf.io/q3zws/

4 http://sccn.ucsd.edu/wiki/Plugin\_list\_process

5 The code is available here: https://github.com/norahollenstein/zuco-benchmark

6 https://scikit-learn.org/stable/modules/generated/sklearn.svm.SVC.html

